# Relative and quantitative rhizosphere microbiome profiling result in distinct abundance patterns

**DOI:** 10.1101/2021.02.19.431941

**Authors:** Hamed Azarbad, Julien Tremblay, Luke D. Bainard, Etienne Yergeau

**Author notes:** **Corresponding author at**: Centre Armand-Frappier Santé Biotechnologie, Institut national de la recherche scientifique, 531 boulevard des Prairies, Laval, QC, H7V 1B7, Canada., **E-mail address**.

## Abstract

Next-generation sequencing is recognized as one of the most popular and cost-effective way of characterizing microbiome in multiple samples. However, most of the currently available amplicon sequencing approaches are inherently limited, as they are often presented based on the relative abundance of microbial taxa, which may not fully represent actual microbiome profiles. Here, we combined amplicon sequencing (16S rRNA gene for bacteria and ITS region for fungi) with real-time quantitative PCR (qPCR) to characterize the rhizosphere microbiome of wheat. We show that the increase in relative abundance of major microbial phyla does not necessarily result in an increase in abundance. One striking observation when comparing relative and quantitative abundances was a substantial increase in the abundance of almost all phyla associated with the rhizosphere of plants grown in soil with no history of water stress as compared with the rhizosphere of plants growing in soil with a history of water stress, which was in contradiction with the trends observed in the relative abundance data. Our results suggest that the estimated absolute abundance approach gives a different perspective than the relative abundance approach, providing complementary information that helps to better understand the rhizosphere microbiome.

## INTRODUCTION

Most of the currently available amplicon sequencing approaches are inherently limited and produce compositional data, often presented as the relative abundance of microbial taxa (fraction of total reads), and are not taking into account inter-sample variations in microbial loads (Props *et al*., 2017). Compositional data is constrained, and the biological interpretation of such datasets can be misleading (Vandeputte *et al*., 2017). This is particularly important when substantial differences in total microbial biomass between samples are expected such as in climate change experiments where stressed samples (e.g. drought, heat, and salinity stresses) are compared to control samples. We know from previous studies that both bacterial and fungal species may show different response patterns in water-limited environments such as sensitive, tolerant and opportunistic (Evans and Wallenstein, 2014; Meisner *et al*., 2018), that ultimately shape community structure and many ecosystem functions. If, for instance, a single opportunistic bacterial species increase in absolute abundance (counts) under water stress, this will result (i) in an increase in its relative abundance (ratios) within the community and (ii) in a decrease in the abundance of all other species due to compositionality effects (Morton *et al*., 2017; Vandeputte *et al*., 2017), which is independent of ecological processes governing community profiles (Stämmler *et al*., 2016).

Here, we compared the usefulness of relative and estimated absolute abundances to assess the responses of wheat rhizosphere microbial communities to long-term water stress. To address this, we examined the rhizosphere microbiome of four wheat genotypes (two recognized for their resistance to water stress and two that are not) growing in soils taken from two semi-arid wheat fields with more than 40 years exposure to different irrigation management histories: one that has been irrigated during the growing season (IR soils) and the other field that was not (NI soils).

## MATERIALS AND METHODS

### Soil sample collections and experimental design

Twenty soil samples were collected in April 2016 from the top layer of two experimental agricultural wheat fields located at the Swift Current Research and Development Centre (Agriculture and Agri-Food Canada) in Swift Current, SK, Canada. Although adjacent, these fields were managed differently since 1981 in such a way that one was irrigated (IR) during the wheat growing season and the other was not (NI). Soil samples were transported to the laboratory, mixed, homogenized, and sieved (2 mm) to obtain a representative soil for each wheat field. Two wheat genotypes with recognized resistance to water stress (*Triticum aestivum* cv. AC Barrie and *Triticum turgidum* subsp. *Durum* cv. Strongfield) and two without (*Triticum aestivum* cv. AC Nass and *Triticum aestivum* cv. AC Walton) were sown in pots containing 700 g of each type of soil (dry weight equivalent) and placed in a growth room. During the first four weeks of the experiment, pots were kept under well-watered conditions (50% soil water holding capacity, SWHC), then they were subjected to 5–8% SWHC, 20% SWHC, and 30% SWHC, while controls were kept at 50% SWHC. Rhizosphere samples were collected (4 wheat genotypes x 2 soil history type x 4 SWHC × 5 replicates = 160 samples) at the end of the experiment (after four weeks of exposure to different soil water content) to measure rhizosphere CO_2_ emission and microbial community structure. Detailed information regarding sampling sites, the experimental design, CO_2_ production measurements, qPCR assays, amplicon library construction and sequencing has been previously published (Azarbad *et al*., 2018, 2020) and is provided as Supplementary Material.

### Statistical analyses

Statistical analyses were carried out using the PAST program (Hammer *et al*., 2001). In this study, absolute gene densities (16S rRNA gene and ITS region qPCR data) varied between the “original soils” before the start of the experiment (that is 3 subsamples of the sieved soil prior to wheat seeding) for both bacteria (IR soils: 3.9 × 10^7^ copies and NI soils: 2.2× 10^7^ copies g^−1^ soil-dw) and fungi (IR soils: 9.8 × 10^6^ copies and NI soils: 1.6 × 10^7^ copies g^−1^ soil-dw). We then measured the number copies of the 16S rRNA gene and of the ITS region in each sample to estimate the absolute abundances of bacteria and fungi and to normalize amplicon sequencing data. Two datasets were produced: OTU relative abundance (fraction of total reads) and absolute OTU abundance which was estimated by multiplying the OTU relative abundance matrix by the corresponding abundance of 16S rRNA gene and ITS region obtained by qPCR quantifications, as previously suggested (Zhang *et al*., 2017; Jian *et al*., 2020). Since soil history was previously identified as the main factor structuring microbial communities (Azarbad *et al*., 2018, 2020), we focused on this factor for the needs of our demonstration, but similar conclusions could be reached by focusing on cultivar or SWC effects. To investigate the possible effect of soil history on the relative and estimated absolute abundance of the most abundant bacterial and fungal phyla associated with the rhizosphere of wheat genotypes, analysis of variance (ANOVA) was performed. To verify which approach is most closely related to the functional processes in the rhizosphere, Pearson correlation tests were performed between the estimated absolute and relative abundances of the dominant bacterial and fungal phyla) and CO_2_ production.

## RESULTS AND DISCUSSION

By comparing the relative and the estimated absolute abundances of dominant bacterial and fungal phyla associated with the rhizosphere, we observed completely different and sometime contradictory trends. For instance, the relative abundances of *Acidobacteria* and *Firmicutes* were significantly higher in the rhizosphere of plants growing in the non-irrigated soil as compared to the irrigated soil (Fig. 1a), whereas the estimated abundances of *Proteobacteria, Actinobacteria, Bacteroidetes, Acidobacteria, Gemmatimonadetes, Firmicutes*, and *Verrucomicrobia* were significantly higher in the rhizosphere of plants grown in irrigated soil when compared to non-irrigated soils (Fig. 1b). In contrast, *Gemmatimonadetes* and *Verrucomicrobia* had significantly higher relative abundances in the rhizosphere of plants grown in irrigated soil as compared to non-irrigated soils, coherent with the picture observed for estimated abundances (Fig. 1a). There were also inconsistent trends between the relative and estimated absolute abundances for fungal phyla (Fig. 2). For instance, the relative abundance of *Zygomycota* and *Ascomycota* increased in the rhizosphere of plants grown in irrigated soils, but this pattern was absent when looking at the estimated abundances (Figs. 2a and b). In contrast, both relative and estimated abundances agreed that a history of water stress significantly increase the abundance of *Basidiomycota* in the rhizosphere of wheat (Figs. 2a and b).

**Figure 1.**
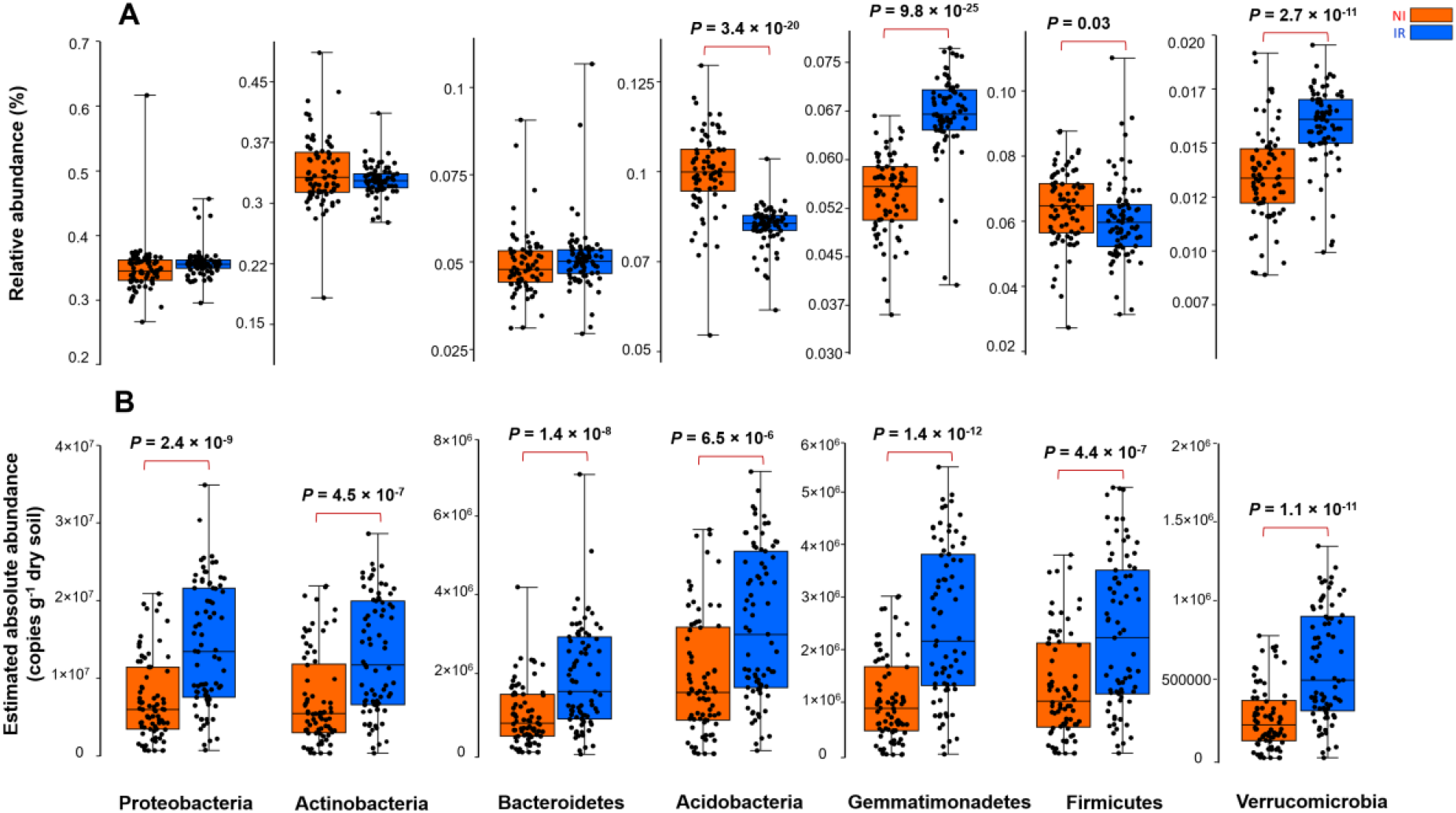
The effect of soil history on (a) the relative and (b) estimated absolute abundance of different bacterial phyla associated with rhizosphere of four wheat genotypes grown in soil with two different water stress history sampled from same wheat field from Saskatchewan with a water stress history (NI) and with no history of water stress (IR) exposed to four level of soil water content. Dots are values for individual observations, the horizontal lines in boxes are representative of median, the upper and lower part of boxes indicating 75th and 25th quartiles, and whiskers on the boxes showing 1.5 × the interquartile range. ANOVA tests comparing the abundance of each phyla between IR and NI soils. ANOVA test, *P < 0.05, **P < 0.01, ***P < 0.001.

**Figure 2.**
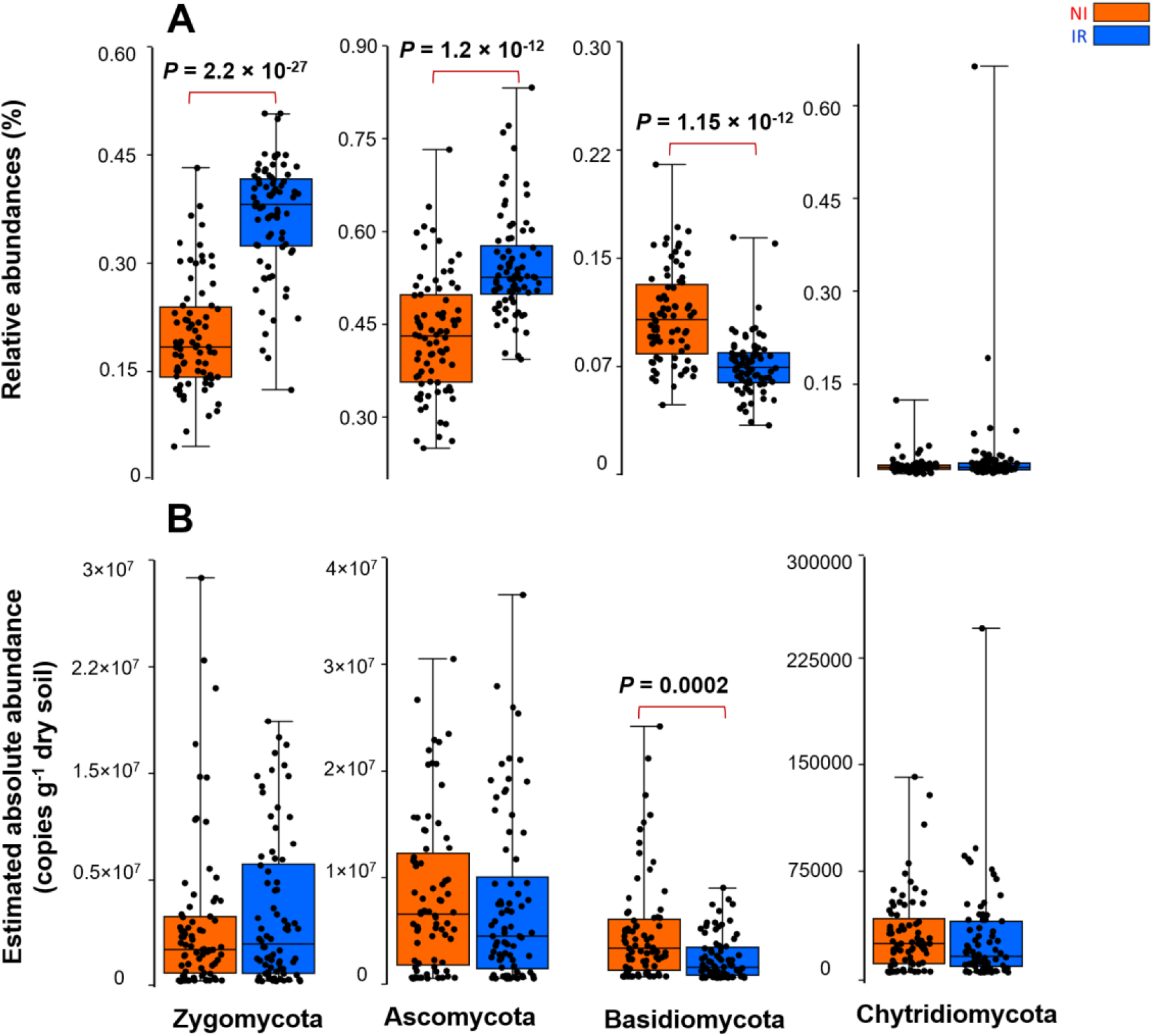
The effect of soil history on (a) the relative and (b) estimated absolute abundance of different fungal phyla associated with rhizosphere of four wheat genotypes grown in soil with two different water stress history sampled from same wheat field from Saskatchewan with a water stress history (NI) and with no history of water stress (IR) exposed to four level of soil water content. Dots are values for individual observations, the horizontal lines in boxes are representative of median, the upper and lower part of boxes indicating 75th and 25th quartiles, and whiskers on the boxes showing 1.5 × the interquartile range. ANOVA tests comparing the abundance of each phyla between IR and NI soils. ANOVA test, *P < 0.05, **P < 0.01, ***P < 0.001.

To determine which approach was more closely linked to the functional processes in the rhizosphere, we correlated relative and absolute abundances of dominant bacterial phyla with CO_2_ production which was previously measured (Azarbad *et al*., 2018). When the estimated absolute abundance was used, Pearson correlations showed no significant correlations between bacterial phyla and CO_2_ production. However, when relative abundance data were used, we observed significant negative (*Actinobacteria*: r = −0.360, p = <0.001) and positive (*Acidobacteria*: r = 0.157, p = 0.048; *Gemmatimonadetes*: r = 0.272, p = <0.001; *Proteobacteria*: r = 0.276, p = <0.001) correlations between bacterial phyla and CO_2_ emissions (Table 1).

**Table 1.** Correlation tests between relative and absolute abundances of the most abundant bacterial phyla associated with rhizosphere of four wheat genotypes grown in soil with two different water stress history sampled from same wheat field from Saskatchewan with a water stress history (NI) and with no history of water stress (IR) exposed to four level of soil water content vs. CO_2_ production.

|  | CO<sub>2</sub> production |  |  |  |
| --- | --- | --- | --- | --- |
|  | Relative abundance |  | Quantitative abundance |  |
|  | R | P Value | R | P Value |
| Acidobacteria | 0.158 | **0.048** | 0.101 | 0.208 |
| Actinobacteria | -0.361 | **<0.001** | 0.040 | 0.615 |
| Bacteroidetes | -0.036 | 0.656 | 0.061 | 0.451 |
| Firmicutes | -0.079 | 0.327 | 0.051 | 0.528 |
| Gemmatimonadetes | 0.272 | **0.001** | 0.104 | 0.193 |
| Proteobacteria | 0.277 | **<0.001** | 0.093 | 0.249 |
| Verrucomicrobia | 0.071 | 0.378 | 0.078 | 0.331 |

In the present study, absolute gene densities (qPCR data) were significantly different between the two soils before starting the experiment, which is one of the many cases for which analyses of microbial communities based on relative abundance is unlikely to reflect actual community patterns (Stämmler *et al*., 2016; Zhang *et al*., 2017; Props *et al*., 2017; Vandeputte *et al*., 2017; Lundberg *et al*., 2020) and associated ecosystem processes. Here, when estimated absolute abundances were considered, we observed notable differences from the relative abundance data and new, often contradictory, trends became visible. It is relatively easy to understand why these differences occur: for example, if a single drought tolerant bacterial species increases its absolute abundance under water stress, this will result in an increase in its relative abundance and to a concomitant decrease in the relative abundance of all other species, even though their absolute abundance did not change. Such compositional effects may not fully reflect actual microbial profiles and their associated ecological functional processes. Using real-time PCR we previously showed that the microbial abundance is quite stable across SWC treatments, for both soil history types (Azarbad *et al*., 2018). However, CO_2_ emissions were severely reduced under low water content, which is clearly due to a change in the activity of the microbial community, rather than a massive death and reduction of abundance of the microbial communities. The estimated absolute data supported this observation as no correlation between abundance of the dominant bacterial phyla and CO_2_ production was detected. Conversely, in the case of the relative abundance approach significant correlations between many taxa and CO_2_ production were found. These correlations were most probably spurious and due to shifts in the microbial community composition with decreasing water content. Some results were even nonsensical, as negative correlations were found, which would be interpreted as if soil respiration would decrease when some taxa become more abundant.

This study demonstrated the potential usefulness of incorporating abundance data derived from widely available qPCR protocols in plant microbiome studies. Since both qPCR and sequencing approaches were performed on the same DNA extract with the same set of primers, they shared the same methodological limitations (e.g. amplification efficiency and the specificity of primers) thereby making the qPCR and sequencing data compatible (Dannemiller *et al*., 2014; Props *et al*., 2017; Jian *et al*., 2020). Alternatively, performing qPCR with taxa-specific primers would provide the ability to track whether a given taxa increased its absolute abundance while other taxa remained the same without the need for sequencing. Other promising approaches such as flow cytometry (Props *et al*., 2017), adding a known number of 16S rRNA gene copies of exogenous bacteria into the samples before DNA extraction in order to normalize endogenous bacterial counts (Stämmler *et al*., 2016) or synthetic spike-in standards (Tourlousse *et al*., 2016; Tkacz *et al*., 2018; Palmer *et al*., 2018) could also be applied to estimate microbial loads.

## Conclusion

In conclusion, this study allowed us to quantitatively evaluate the differences between the rhizosphere microbiome of wheat plants growing in soil with contrasting long-term water management history. We showed that quantitative microbiome profiling provides a contrasting picture of the response of rhizosphere microbial communities to soil water stress legacy, which appeared to be better aligned to actual ecosystem processes.

## Acknowledgements

This work was funded by the Natural Sciences and Engineering Research Council of Canada (Discovery grant RGPIN-2014-05274 and Strategic grant for projects STPGP 494702 to EY). HA was supported by a Fonds de recherche du Québec-Nature et technologies (FRQNT) and a Fondation Armand-Frappier postdoctoral fellowships. This research was enabled in part by support provided by Calcul Québec (www.calculquebec.ca) and Compute Canada on the Graham compute cluster (University of Waterloo) (www.computecanada.ca).

