## Supplementary Methods for "Relative and quantitative rhizosphere microbiome profiling result in distinct abundance patterns"

*DNA extraction and qPCR assays*

Genomic DNA was extracted from 0.5 mg of rhizosphere samples using a phenol-chloroform extraction method (Dellaporta *et al*., 1983). Further details on DNA extraction can be found in Azarbad *et al*. (2018). Total abundance of bacterial (16S rRNA genes) and fungal (ITS1 region) communities associated with the rhizosphere, root and leaf were quantified using SyBrGreen real-time quantitative PCR assays (qPCR) with the primers listed below (same as those used for amplicon sequencing assays). Briefly, the qPCR reactions were carried out using RotorGene 6000 machine (Corbett Research, Mortlake, NSW, Australia) with SsoAdvanced™ Universal SYBR Green kit (Biorad, Hercules, CA, USA). We have performed several tests runs including wide range of dilution of extracted DNA in order to: (1) find out the range of linear amplification of extracted DNA to ensure that all samples are in expected scale based on the standard curve and (2) to reduce qPCR inhibition. The standard curve ranging from 0 to $10^7$ copies of the standard plasmid DNA were prepared using *Escherichia coli* 25922 for bacteria (Bruce *et al*., 1992) and *Pichia scolyti* for fungi (Martin and Rygiewicz, 2005). We have chosen 10 random samples and then prepared 10, 50, 100, 200 and 400-fold diluted and non-diluted DNA extracts. We also included two blank samples (nuclease-free water) as controls. Based on these results, the optimum dilutions were selected if the amplification products were between expected ranges (above the minimum detection limit to the middle point of the linear range of standards). As a result, DNA fragments corresponding to the rhizosphere were diluted 10 times, respectively. Each qPCR mix consisted of 4.2 µl sterilized water, 10 µl SYBR green master mix, 0.4 µl of each primer (0,4 pmoles/µl) and 5 µl (of diluted) template DNA were used to have a final reaction volume of 20 µl. Each qPCR run had at least two no template controls. The PCR conditions consists of an initial denaturation step at 95°C for 5 minutes followed by 30 cycles of denaturation at 95°C for 30 s, annealing at 57°C for 30 s and elongation at 72°C for 30 s. Fluorescence was measured at the end of each cycle at the elongation step. A melt curve analysis was done to verify the specificity of the amplicons.

*Amplicon library construction and sequencing*

Sequencing libraries were created as described previously in Yergeau *et al*. (2015) based on dual-indexed strategy following the “16S Metagenomic Sequencing Library preparation” Illumina guide (Part #15044223 Rev. B). Similar to qPCR analysis, for bacterial 16S rRNA gene the V3-V4 hypervariable region was amplified using the universal primers 520F (5’-AGCAGCCGCGGTAAT-3’) and 799R (5’-CAGGGTATCTAATCCTGTT-3’) (Edwards *et al*., 2008) and for fungi the ITS1 region was amplified using ITS1F (5’-CTTGGTCATTTAGAGGAAGTAA-3’) and 58A2R (5’-CTGCGTTCTTCATCGAT-3’) (Martin and Rygiewicz, 2005). Samples were pooled separately for fungi and bacteria and submitted for 2 × 250 bp Illumina MiSeq sequencing at the McGill University and Genome Québec Innovation Centre (Montréal, Canada). Sequence data were analysed following procedures described in Tremblay *et al*. (2015). Briefly, raw reads were controlled for quality. Remaining high quality reads and free of sequencing adapters artefacts were dereplicated at 100% identity and clustered/denoised at 99% (DNAclust v3) (PMID:21718538). Clusters of less than three reads were discarded and remaining clusters were scanned for chimeras using UCHIME, first in de novo mode then in reference mode (Edgar *et al*., 2011). Remaining clusters were clustered at 97% identity (DNAclust v3) to produce OTUs. For 16S data types, taxonomy assignment of resulting OTUs was performed using the RDP classifier (Wang *et al*., 2007) with a modified Greengenes training set built from a concatenation of the Greengenes database v13_5 (DeSantis *et al*., 2006), and Silva eukaryotes 18S r128 (Quast *et al*., 2013). For ITS data, taxonomic assignment was done with the RDP classifier using a training set generated from the Unite database (sh_refs_qiime_ver7_dynamic_20.11.2016) (Kõljalg *et al*., 2013). Raw data sets are available in the NCBI Sequence Read Archive (SRA) under the BioProject accession PRJNA526458.
